# M-type potassium currents differentially affect activation of motoneuron subtypes and tune recruitment gain

**DOI:** 10.1101/2023.07.24.550111

**Authors:** Simon A. Sharples, Matthew J. Broadhead, James A. Gray, Gareth B. Miles

## Abstract

The size principle is a key mechanism governing the orderly recruitment of motor units and is believed to be dependent on passive properties of the constituent motoneurons. However, motoneurons are endowed with voltage-sensitive ion channels that create non-linearities in their input-output functions. Here we describe a role for the M-type potassium current, conducted by KCNQ channels, in the control of motoneuron recruitment in mice. Motoneurons were studied with whole-cell patch clamp electrophysiology in transverse spinal slices and identified based on delayed (fast) and immediate (slow) onsets of repetitive firing. M-currents were larger in delayed compared to immediate firing motoneurons, which was not reflected by variations in the expression of Kv7.2 or Kv7.3 subunits. Instead, a more depolarized spike threshold in delayed-firing motoneurons afforded a greater proportion of the total M-current to become activated within the subthreshold voltage range, which translated to a greater influence on their recruitment with little influence on their firing rates. Pharmacological activation of M-currents also influenced motoneuron recruitment at the population level, producing a rightward shift in the recruitment curve of monosynaptic reflexes within isolated mouse spinal cords. These results demonstrate a prominent role for M-type potassium currents in regulating the function of motor units, which occurs primarily through the differential control of motoneuron subtype recruitment. More generally, these findings highlight the importance of active properties mediated by voltage-sensitive ion channels in the differential control of motoneuron recruitment, which is a key mechanism for the gradation of muscle force.

**Key Points:**

- M-currents exert an inhibitory influence on spinal motor output.
- This inhibitory influence is exerted by controlling the recruitment, but not the firing rate, of high-threshold fast-like motoneurons, with limited influence on low-threshold slow-like motoneurons.
- Preferential control of fast motoneurons may be linked to a larger M-current that is activated within the subthreshold voltage range compared to slow motoneurons.
- Larger M-currents in fast compared to slow motoneurons are not accounted for by differences in Kv7.2 or Kv7.3 channel composition.
- The orderly recruitment of motoneuron subtypes is shaped by differences in the contribution of voltage-gated ion channels, including KCNQ channels.
- KCNQ channels may provide a target to dynamically modulate the recruitment gain across the motor pool and readily adjust movement vigour.

## Introduction

Complex behaviour is critically dependent on the ability to readily adjust the degree of precision and vigour of movement. This adaptability in motor control relies on gain control strategies, which include the orderly recruitment and subsequent firing rate modulation of motor units; the smallest functional unit of the motor system composed of a motoneuron and the muscle fibres it innervates. It is well established that the orderly recruitment of motor units follows the size principle, with sequential recruitment of motor units of increasing size (Henneman, 1957). Motor units can also be classified based on muscle twitch properties, with sequential recruitment of motor units that produce greater muscle force, greater speed, and exhibit increased susceptibility to fatigue (Burke *et al*., 1973). However, in contrast to what would be predicted based on the size principle, motoneuron size is a poor predictor of recruitment within functionally-defined motor unit subtypes (Gustafsson & Pinter, 1984; Sharples & Miles, 2021). Further, motoneuron size is a property that is not readily adjustable on the timescale that is needed to produce flexible output. Emerging evidence points toward a role for active properties mediated by voltage-gated ion channels in shaping motoneuron recruitment, with differences in their expression and voltage gating affording differential control of motoneuron subtypes (Sharples & Miles, 2021). Differences in the complements of voltage-gated ion channels are not only important to support the physiological functions of different types of motoneurons, but may also underlie differential susceptibility to degeneration in a range of disorders of the nervous system (Rose & Griggs, 2001; Nijssen *et al*., 2017; Ragagnin *et al*., 2019; Soulard *et al*., 2020).

The M-type potassium current, conducted by KCNQ channels, is a voltage-dependent, non–inactivating outward current that is activated within the subthreshold voltage range and plays a key role in the control of neuronal excitability across the central nervous system (CNS). The M-current has been implicated in a range of neural disorders including epilepsy, neuropathic pain, and Amyotrophic Lateral Sclerosis, which are characterized by aberrant neuronal excitability leading to excitotoxicity and cell death (Blackburn-Munro & Jensen, 2003; Gunthorpe *et al*., 2012; Wainger *et al*., 2014, 2021). As a result, these channels have become a key therapeutic target to combat disease progression (Wainger *et al*., 2014, 2021; Buskila *et al*., 2019). In the spinal cord, M-currents are found in excitatory, rhythm generating spinal interneurons where they play a key role in producing the underlying rhythms that are important for locomotor movement (Verneuil *et al*., 2020). M-currents have also been found in lumbar motoneurons in turtles (Alaburda *et al*., 2002) and hypoglossal motoneurons in rats (Ghezzi *et al*., 2017, 2018). While KCNQ channels are expressed in lumbar motoneurons of mice (Verneuil *et al*., 2020), it is not yet known how these channels might provide differential control motoneuron subtypes.

Here, we present a novel role for M-type potassium currents in the gain control of spinal motor output. Using whole-cell patch clamp electrophysiology, we show that M-currents are larger in higher threshold fast subtypes of motoneurons, which is not reflected in differences in Kv 7.2 or Kv7.3 subunit expression. Pharmacological manipulation of M-currents modulates the recruitment, but not firing rates of fast but not slow type motoneurons. This finding was recapitulated at the population level as a right shift in monosynaptic reflex recruitment curves in isolated mouse spinal cords following pharmacological activation of KCNQ channels. Together, these data advance our understanding of gain control mechanisms in the motor system, describing novel roles for ion channels in the neural control of movement.

## Results

### Differential expression of M-type potassium currents in motoneuron subtypes

Whole-cell patch clamp electrophysiology was deployed to study the intrinsic properties of motoneurons in transverse lumbar slices that were obtained from mice during the second two postnatal weeks (P10-18). In these preparations, correlates of fast and slow motoneuron subtypes can be identified based on a delayed or immediate onset of repetitive firing during a 5 second depolarizing current step (Figure 1A, B; (Leroy *et al*., 2014, 2015; Bos *et al*., 2018; Sharples & Miles, 2021)). We first set out to determine whether we could identify putative M-currents in delayed and immediate firing motoneurons. Putative M-currents were studied in voltage clamp using an established deactivation protocol whereby the membrane potential was hyperpolarized from -10 mV, which is a voltage where M-currents would be expected to be maximally activated. Motoneurons were then subjected to 1-second hyperpolarizing voltage steps of 5 mV increments, which produced a negative current deflection as a result of M current deactivation (Figure 1C, D; (Adams *et al*., 1982; Ghezzi *et al*., 2017; Verneuil *et al*., 2020). Blockade of KCNQ channels (XE991; 10 µM) led to a significant reduction in maximal current in both delayed (- 50 ± 24 %) and immediate (- 62 ± 22 %) firing motoneurons (Figure 1E - J)**^a^**, suggesting that both motoneuron types do indeed express M-currents.

**Figure 1:**
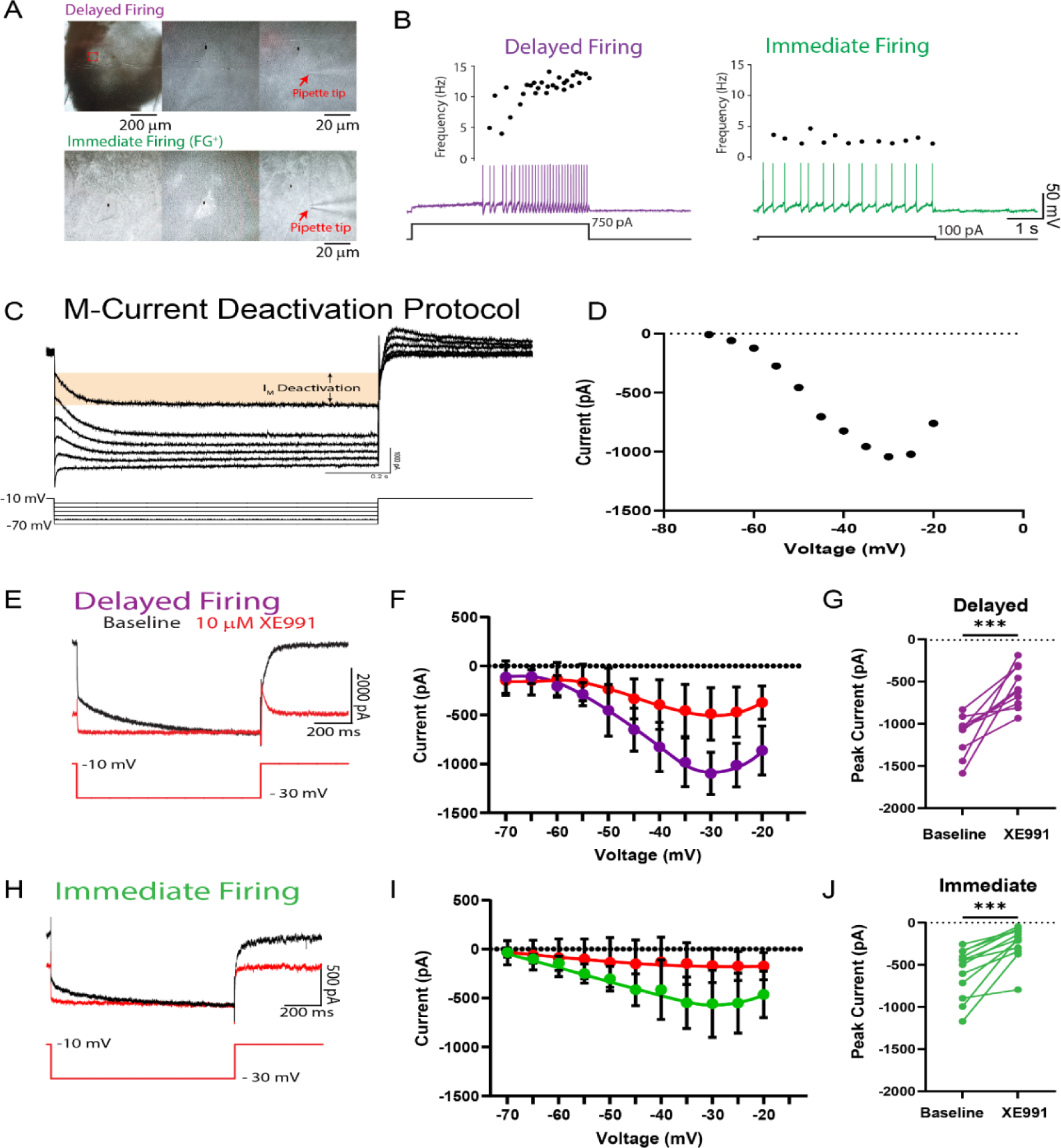
Delayed and immediate firing MNs express an XE991-sensitive M-current. (A) Motoneurons visualized under differential interference contrast were identified based on delayed (purple) and immediate (green) firing with long (5 second) depolarizing current steps applied near rheobase current (B). (C) Putative M-type potassium currents were studied using a deactivation protocol. (D) Current-Voltage relationship for putative M-type potassium current. Putative M-type potassium currents recorded in delayed (E-G) and immediate (H-J) firing motoneurons could be blocked by the KCNQ channel blocker XE991 (10 µM; n = 9 delayed; n = 12 immediate). Data in F and I are presented as mean ± SD. Data in G and J were analyzed with a paired t-test. Asterisks denote significant differences * p<0.05, *** p<0.01; *** p <0.001.

Having identified M-currents in delayed and immediate firing motoneurons, we next set out to determine whether M-currents differed between motoneuron subtypes (Figure 2A). M-currents were significantly larger in delayed compared to immediate firing motoneurons (Figure 2B)**^b^**, with no differences in current half deactivation voltage (Figure 2C)**^c^**, or deactivation time constant (delayed: 125 ± 82 ms, Immediate: 133 ± 73 ms)**^d^**. M-current amplitude did not change across the second and third postnatal week, however differences between subtypes were preserved**^e^**.

**Figure 2:**
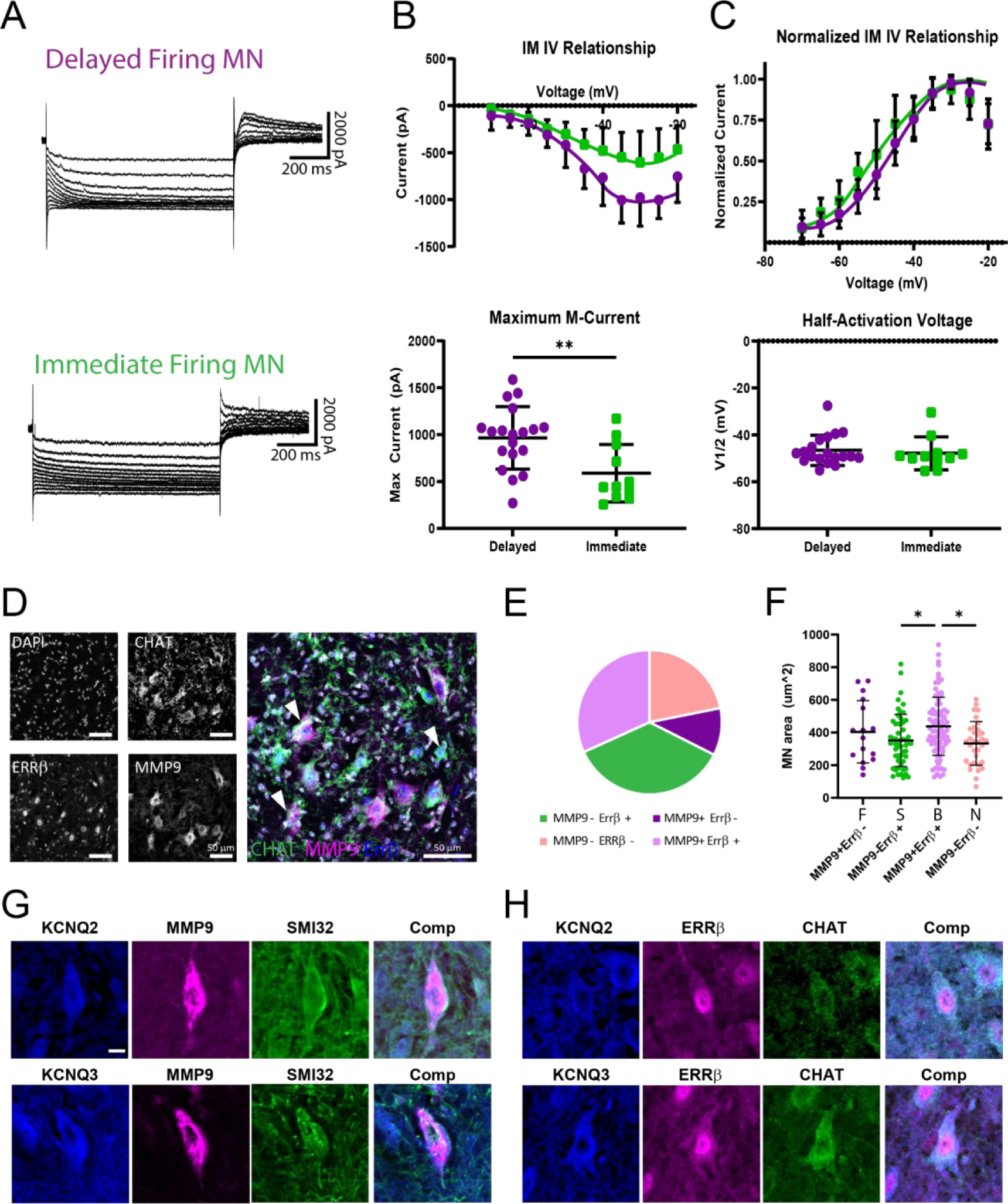
M-currents are larger in delayed compared to immediate firing motoneurons. (A) Representative traces of M-currents measured in 18 delayed (purple) and 15 immediate firing motoneurons (green) (B) Current-voltage relationship for M-type potassium currents in delayed (purple) and immediate (green) firing motoneurons. Maximal M-currents were significantly larger in delayed firing motoneurons. (C) Normalized current voltage relationship revealed no differences in half-activation voltage of M-currents between subtypes. Data are presented as mean ± SD, and individual data points shown. Data were analyzed with 2 factor repeated measures ANOVA (IV plots) or unpaired t-tests. Asterisks denote statistical significance with **p<0.01. (D) Immunohistochemical labelling for DAPI (White), Choline Acetyl Transferase (ChAT; Green); Matrix Metallopeptidase 9 (MMP9; Magenta), and the Estrogen-Related Receptor Beta (Errβ; Blue) reveal heterogeneity among molecularly defined motoneuron subtypes. (E) Pie graph illustrating proportion of ChAT+ motoneurons that express combinations of MMP9 and Errβ. (F) Motoneurons that express MMP9 are the largest, regardless of presence of Errβ. (G) Fast fatigable motoneurons co-labelled for MMP9 (magenta) and SMI-32 (green) express both KCNQ2 and KCNQ3 channel subunits (blue). (H) Slow motoneurons co-labelled for Errβ (magenta) and ChAT (green) express both KCNQ2 and KCNQ3 channel subunits (blue). Scale bar 10 um.

M-currents are conducted by heteromeric KCNQ channels composed of five Kv channel subunits (Kv 7.1 – 7.5). Previous work has demonstrated that all spinal motoneurons express Kv7.2 subunits, with only 50% expressing Kv 7.3 (Verneuil *et al*., 2020). Given that Kv 7.2 and Kv7.3 heteromers can lead to larger M-currents (Jentsch, *et al.,* 2000), we hypothesized that fast motoneurons would express both Kv7.2 and Kv7.3 subunits, whereas the slow motoneurons would express only Kv 7.2 subunits. To address this question, we performed immunohistochemical labelling for Kv7.2 and Kv7.3 channels in putative fast fatigable and slow type motoneurons identified by their expression of cytoplasmic matrix metalloproteinase 9 (MMP9) (Kaplan *et al*., 2014; Leroy *et al*., 2014; Martínez-Silva *et al*., 2018; Allodi *et al*., 2021) or nuclear estrogen-related receptor Beta (Errβ) (Enjin *et al*., 2010; Leroy *et al*., 2014) respectively. Immunohistochemical labelling for choline-acetyltransferase (ChAT) in lamina IX revealed a mixed distribution of cells that expressed MMP9 (10 %), Errβ (36 %), or neither (22 %) (n = 150 MNs, N = 9 section; N = 3 animals). Interestingly, we also found a large proportion (32 %) of motoneurons that express both MMP9 and Errβ (Figure 2D, E), with MMP9-expressing cells being the largest regardless of whether they expressed Errβ or not (Figure 2F). Subsequent analysis of Kv7.2 and Kv7.3 revealed the presence of both subunits in fast motoneurons that were co-labelled with MMP9^+^ and SMI-32 or slow motoneurons co-labelled with Errβ and ChAT, suggesting that overt differences in channel composition does not contribute to larger currents in fast motoneuron subtypes.

### M-type potassium currents control the rheobase of delayed firing motoneurons

In addition to having larger M-currents, delayed firing motoneurons also have a more depolarized spike threshold compared to immediate firing motoneurons (Figure 3A, B)**^f^**, which allows a greater proportion of their M-current to become activated within the subthreshold voltage range (Delayed: 80 ± 0.2 %; Immediate: 52 ± 0.25 %; Figure 4C)**^g^**. We therefore hypothesized that differences in the amount of M-current activated below spike threshold would translate to a greater influence on their recruitment. In line with our hypothesis, the M-current blocker XE991 (10 µM) decreased the rheobase of delayed (Figure 3D) but not immediate firing (Figure 3E) motoneurons, to a level where rheobase current of delayed firing motoneurons was not significantly different (p = 0.2) from that of immediate firing motoneurons (Figure 3F)**^h^**. This nonuniform modulation of rheobase currents across motoneuron subtypes led to a 49 % reduction in the rheobase current range (BL: 1873 pA; XE991: 950 pA) and an increase in the recruitment gain across the sample of motoneurons that we studied (Figure 3G). The reduction in rheobase of delayed firing motoneurons was paralleled by a significant increase in the input resistance of both motoneuron subtypes (Figure 3H)**^h^**, with no change in spike threshold (Figure 3I)**^j^**. XE991 did not alter resting membrane potential **^k^** or the amount of bias current needed to hold cells at -60 mV **^l^**in either motoneuron subtype (Table 1), suggesting that M-currents do not contribute to the resting potential in either subtype.

**Figure 3:**
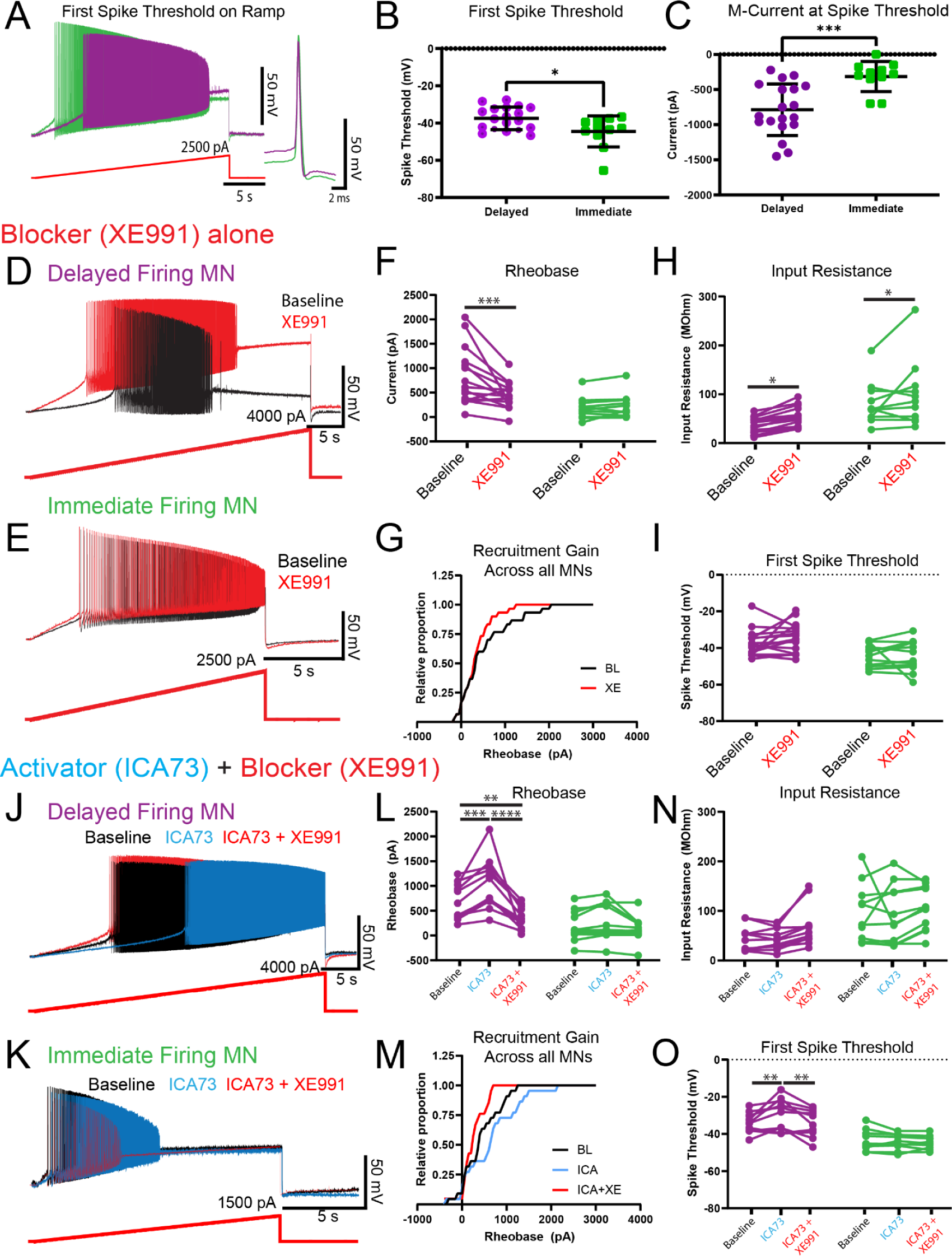
M-currents differentially control recruitment of motoneuron subtypes. (A) Voltage traces from a delayed (purple) and immediate firing (green) motoneuron during slow depolarizing current ramps (red). The first action potential generated at rheobase illustrates differences in spike threshold between the two subtypes. The threshold of the first action potential on depolarizing current ramps is more depolarized in delayed compared to immediate firing motoneurons (B), allowing for a greater amount of M-current to be activated below spike threshold in delayed firing motoneurons (C).Voltage traces from delayed firing (D; n = 16) and immediate firing (E; n = 13) motoneurons before (black) and after (red) application of the KCNQ channel blocker XE991 (10 µM). (F) XE991 significantly decreased the rheobase of delayed firing motoneurons, which translated to a left shift in the recruitment gain across all motoneurons (G). XE991 increased the input resistance of both motoneuron types (H) but did not alter their spike threshold voltage (I). Voltage traces from delayed firing (J; n = 11) and immediate firing (K; n = 11) motoneurons before (black) and after (blue) application of the KCNQ channel activator ICA73 (10 µM). ICA73 increased the rheobase of delayed but not immediate firing motoneurons (L), which led to a right shift in the recruitment gain of all motoneurons (M). ICA73 did not alter the input resistance of either motoneuron subtype (N) but depolarized the threshold of the first action potential on current ramps in delayed firing motoneurons only (O). These effects were reversed with subsequent application of the KCNQ channel blocker, XE991 (10 µM; red). Data are presented as individual points for each motoneuron and analyzed with a 2 factor repeated measures ANOVA. Holm Sidak post hoc analyses were performed when significant differences were found * p<0.05, ** p<0.01; *** p <0.001; **** p < 0.0001.

**Figure 4:**
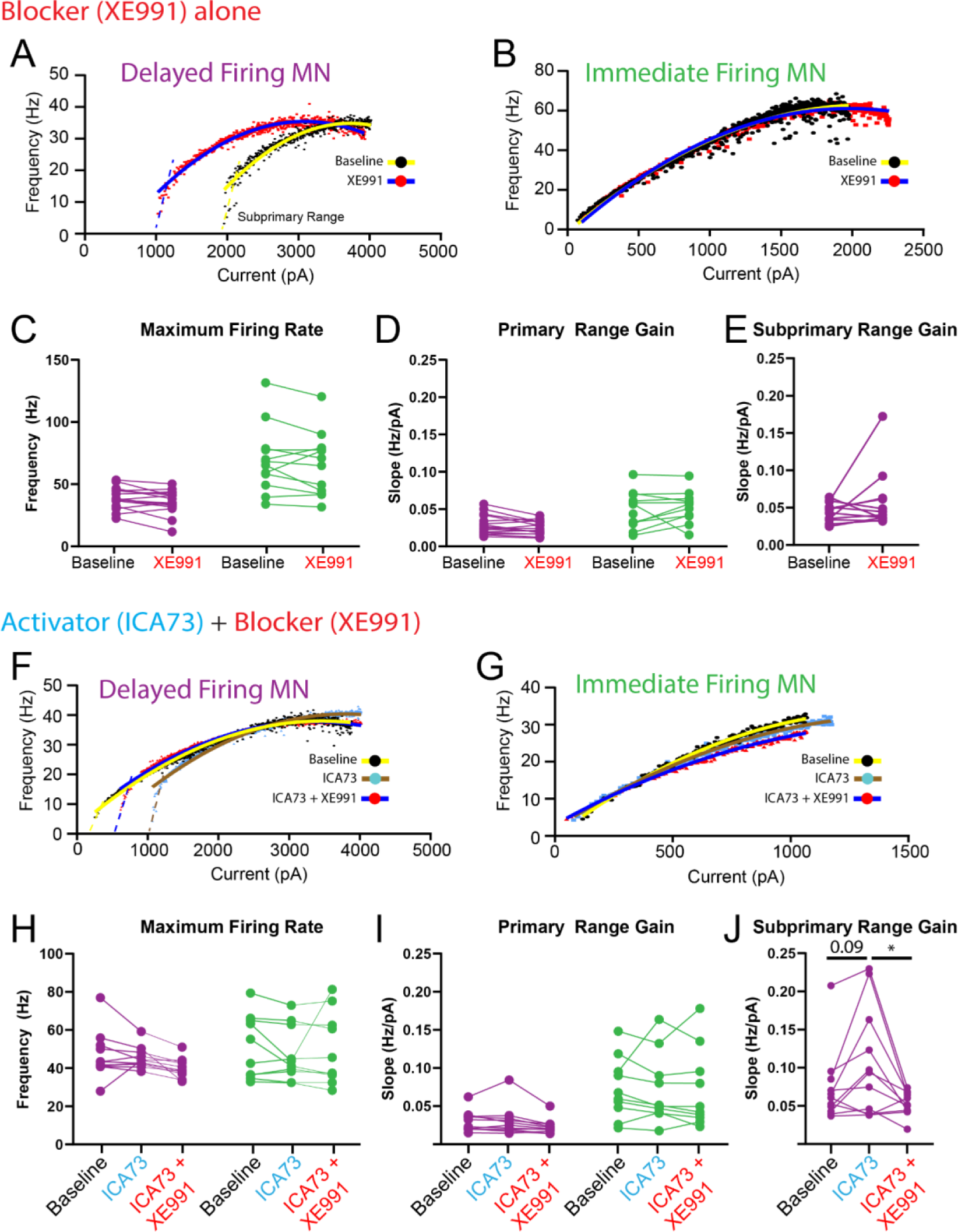
M-currents exert a limited influence on motoneuron firing rate. Frequency-current (f-I) relationship of delayed (A; n = 16) and immediate (B; n = 13) firing motoneurons during slow depolarizing current ramps before (black dots) and after application of the KCNQ channel blocker XE991 (10 µM; red dots). The primary firing range of the f-I relationship in both subtypes can be captured with a second order polynomial (solid lines) and maximal firing rate reached within the plateau region. Delayed but not immediate firing motoneurons also display a linear sub-primary firing range (dashed lines). XE991 produced a left shift in the f-I relationship of delayed firing motoneurons only, but did not affect the maximal firing rate (C) or primary firing range gain (D) of either motoneuron type, nor did it alter the gain within the sub-primary firing range in delayed firing motoneurons (E). Activation of KCNQ channels with ICA73 (10uM: blue dots) produced a right shift in the f-I relationship in delayed firing motoneurons (F; n = 11) but did not alter the f-I relationship of immediate firing (G; n = 11) motoneurons. Further, ICA73 did not change the maximum firing rate (H) or the gain of the primary firing range of either subtype (I). ICA73 increased the gain within the sub-primary firing range of delayed firing motoneurons; an effect that was reversed with subsequent application of the KCNQ channel blocker XE991 (J). Data are presented as individual data points and analyzed with a 2 factor repeated measures ANOVA. Holm Sidak post hoc analyses were performed when significant differences were found * p<0.05.

**Table 1:**
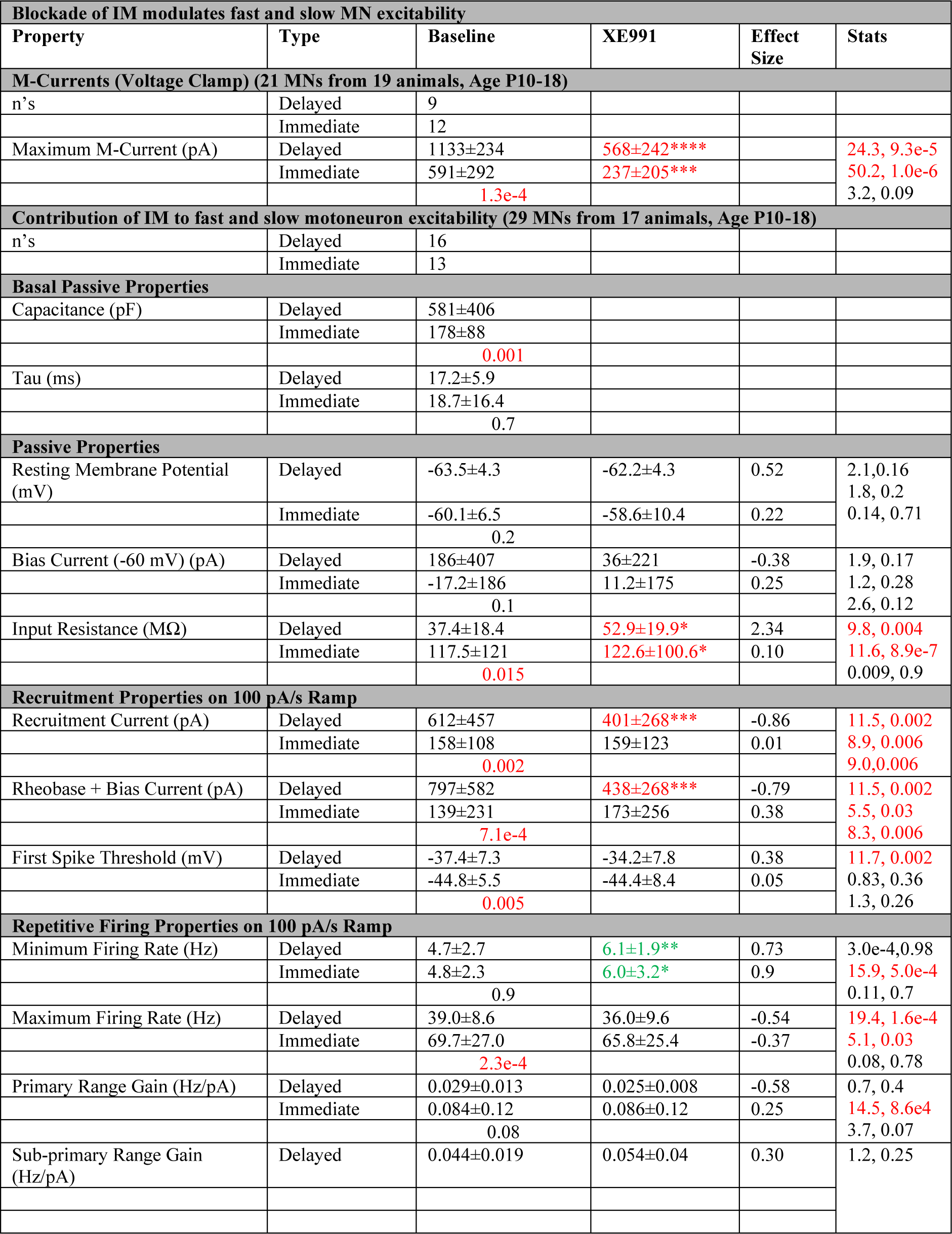
Effects of blocking KCNQ channels on intrinsic properties of motoneuron subtypes. Data are presented as mean ± SD. Asterisks denote significant difference from baseline. F and p values from 2-way Repeated Measures ANOVA for Main effect of MN type, and Main effect of drug, interaction (MN type x Drug). Effect size of antagonist: Small: 0.2 – 0.5; medium: 0. 5- 0.8; Large: > 0.8. Asterisks represent differences from baseline p*p<0.05, **p<0.01, ***p<0.001, ****<0.0001.

| Blockade of IM modulates fast and slow MN excitability |  |  |  |  |  |
| --- | --- | --- | --- | --- | --- |
| Property | Type | Baseline | XE991 | Effect Size | Stats |
| **M-Currents (Voltage Clamp) (21 MNs from 19 animals, Age P10-18)** |  |  |  |  |  |
| n's | Delayed | 9 |  |  |  |
|  | Immediate | 12 |  |  |  |
| Maximum M-Current (pA) | Delayed | 1133±234 | 568±242**** | | 24.3, 9.3e-5 |
| | Immediate | 591±292 | 237±205*** | | 50.2, 1.0e-6 |
|  |  | 1.3e-4 |  |  | 3.2, 0.09 |
| **Contribution of IM to fast and slow motoneuron excitability (29 MNs from 17 animals, Age P10-18)** |  |  |  |  |  |
| n's | Delayed | 16 |  |  |  |
|  | Immediate | 13 |  |  |  |
| **Basal Passive Properties** |  |  |  |  |  |
| Capacitance (pF) | Delayed | 581±406 | | | |
| | Immediate | 178±88 | | | |
|  |  | 0.001 |  |  |  |
| Tau (ms) | Delayed | 17.2±5.9 | | | |
| | Immediate | 18.7±16.4 | | | |
|  |  | 0.7 |  |  |  |
| **Passive Properties** |  |  |  |  |  |
| Resting Membrane Potential (mV) | Delayed | -63.5±4.3 | -62.2±4.3 | 0.52 | 2.1, 0.16 |
| | Immediate | -60.1±6.5 | -58.6±10.4 | 0.22 | 1.8, 0.2 |
|  |  | 0.2 |  |  | 0.14, 0.71 |
| Bias Current (-60 mV) (pA) | Delayed | 186±407 | 36±221 | -0.38 | 1.9, 0.17 |
| | Immediate | -17.2±186 | 11.2±175 | 0.25 | 1.2, 0.28 |
|  |  | 0.1 |  |  | 2.6, 0.12 |
| Input Resistance (MΩ) | Delayed | 37.4±18.4 | 52.9±19.9* | 2.34 | 9.8, 0.004 |
| | Immediate | 117.5±121 | 122.6±100.6* | 0.10 | 11.6, 8.9e-7 |
|  |  | 0.015 |  |  | 0.009, 0.9 |
| **Recruitment Properties on 100 pA/s Ramp** |  |  |  |  |  |
| Recruitment Current (pA) | Delayed | 612±457 | 401±268*** | -0.86 | 11.5, 0.002 |
| | Immediate | 158±108 | 159±123 | 0.01 | 8.9, 0.006 |
|  |  | 0.002 |  |  | 9.0, 0.006 |
| Rheobase + Bias Current (pA) | Delayed | 797±582 | 438±268*** | -0.79 | 11.5, 0.002 |
| | Immediate | 139±231 | 173±256 | 0.38 | 5.5, 0.03 |
|  |  | 7.1e-4 |  |  | 8.3, 0.006 |
| First Spike Threshold (mV) | Delayed | -37.4±7.3 | -34.2±7.8 | 0.38 | 11.7, 0.002 |
| | Immediate | -44.8±5.5 | -44.4±8.4 | 0.05 | 0.83, 0.36 |
|  |  | 0.005 |  |  | 1.3, 0.26 |
| **Repetitive Firing Properties on 100 pA/s Ramp** |  |  |  |  |  |
| Minimum Firing Rate (Hz) | Delayed | 4.7±2.7 | 6.1±1.9** | 0.73 | 3.0e-4, 0.98 |
| | Immediate | 4.8±2.3 | 6.0±3.2* | 0.9 | 15.9, 5.0e-4 |
|  |  | 0.9 |  |  | 0.11, 0.7 |
| Maximum Firing Rate (Hz) | Delayed | 39.0±8.6 | 36.0±9.6 | -0.54 | 19.4, 1.6e-4 |
| | Immediate | 69.7±27.0 | 65.8±25.4 | -0.37 | 5.1, 0.03 |
|  |  | 2.3e-4 |  |  | 0.08, 0.78 |
| Primary Range Gain (Hz/pA) | Delayed | 0.029±0.013 | 0.025±0.008 | -0.58 | 0.7, 0.4 |
| | Immediate | 0.084±0.12 | 0.086±0.12 | 0.25 | 14.5, 8.6e4 |
|  |  | 0.08 |  |  | 3.7, 0.07 |
| Sub-primary Range Gain (Hz/pA) | Delayed | 0.044±0.019 | 0.054±0.04 | 0.30 | 1.2, 0.25 |

In our next set of experiments, we examined how activation of KCNQ channels influences excitability of motoneuron subtypes using the M-current activator ICA73 (10 µM); which hyperpolarizes the activation voltage for KCNQ channels (Verneuil et al., 2020). ICA73 increased the rheobase of delayed (Figure 3J) but not immediate (Figure 3K) firing motoneurons (Figure 3L)**^m^**, producing a 109 % increase in the rheobase current range (BL: 997 pA; ICA73:2082 pA) and a decrease in the recruitment gain across the motor pool (Figure 3M). This decrease in rheobase was not paralleled by a change in input resistance (Figure 3N)**^n^**, but was associated with a depolarization of the spike threshold in delayed firing motoneurons (Figure 3O)**°**. Interestingly, ICA73 significantly hyperpolarized the resting membrane potential (Table 2)**^p^** of delayed firing motoneurons indicating that M-currents do have the capacity to regulate the resting potential. The increase in rheobase, decrease in recruitment gain, depolarization of spike threshold, and hyperpolarization of the resting potential produced by ICA73 in delayed firing motoneurons was reversed by subsequent application of XE991 (Figure 3L-O; Table 2) and was also reproduced in a separate set of experiments by a second M-current activator (Retigabine; 10 µM; n = 7 delayed, n = 4 immediate; Table 3)^q-s^.

**Table 2:** Effects of activating KCNQ channels on intrinsic properties of motoneuron subtypes. Data are presented as mean ± SD. Asterisks denote significant difference from baseline. F and p values from 2-way Repeated Measures ANOVA for Main effect of MN type, and Main effect of drug, interaction (MN type x Drug). Effect size of agonist: Small: 0.2 – 0.5; medium: 0. 5- 0.8; Large: > 0.8. Asterisks represent differences from baseline p*p<0.05, **p<0.01, ***p<0.001, ****<0.0001.

| Activation of IM to modulates fast and slow motoneuron excitability (22 MNs from 14 animals, Age P10-16) |  |  |  |  |  |  |
| --- | --- | --- | --- | --- | --- | --- |
| Property | Type | Baseline | ICA73 | ICA73+XE991 | Effect Size | Stats (F, p) |
| n's | Delayed | 11 |  |  |  |  |
|  | Immediate | 11 |  |  |  |  |
| **Basal Passive Properties** |  |  |  |  |  |  |
| Capacitance (pF) | Delayed | 479±263 | | | | |
| | Immediate | 213±125 | | | | |
|  |  | 0.009 |  |  |  |  |
| Tau (ms) | Delayed | 15.8±4.2 | | | | |
| | Immediate | 15.0±8.2 | | | | |
|  |  | 0.8 |  |  |  |  |
| **Passive Properties** |  |  |  |  |  |  |
| Resting Membrane Potential (mV) | Delayed | -63±2.4 | -66±3.6** | -61±3.6* | -1.0 | 0.2, 0.65 |
| | Immediate | -63±6.2 | -65±6.3 | -64±4.6 | -0.99 | 11.3, 0.02 |
|  |  | 0.99 |  |  |  | 4.4, 1.4e-4 |
| Bias Current (-60 mV) (pA) | Delayed | 128±136 | 306±170** | 58±77 | 0.86 | 3.9, 0.06 |
| | Immediate | 1.0±268 | 61±255 | -15±167 | 0.60 | 17.2, 5.5e-6 |
|  |  | 0.2 |  |  |  | 3.7, 0.03 |
| Input Resistance (MΩ) | Delayed | 43.8±25.2 | 42.7±20.6 | 68.4±40.9 | -0.10 | 7.3, 0.01 |
| | Immediate | 93.5±58.5 | 91.6±58.8 | 106±44.2 | -0.03 | 4.7, 0.01 |
|  |  | 0.02 |  |  |  | 0.44, 0.65 |
| **Recruitment Properties on 100 pA/s Ramp** |  |  |  |  |  |  |
| Rheobase (pA) | Delayed | 563±309 | 789±527** | 404±272* | 0.75 | 12.4, 0.002 |
| | Immediate | 188±119 | 203±178 | 148±163 | 0.10 | 13.3, 3.7e-5 |
|  |  | 0.001 |  |  |  | 57.7, 0.001 |
| Rheobase + Bias Current (pA) | Delayed | 691±359 | 1067±524*** | 366±231** | 1.31 | 15.1, 0.009 |
| | Immediate | 189±307 | 264±368 | 134±254 | 0.76 | 21.7, 4.2e-7 |
|  |  | 0.002 |  |  |  | 10.2, 0.003 |
| First Spike Threshold (mV) | Delayed | -33.8±5.2 | -29.0±8.0** | -33.0±7.8 | 0.83 | 22.2, 0.0002 |
| | Immediate | -43.6±5.7 | -44.4±4.2 | -44.6±3.5 | -0.33 | 4.8, 0.01 |
|  |  | 6.3e-4 |  |  |  | 6.1, 0.005 |
| **Repetitive Firing Properties on 100 pA/s Ramp** |  |  |  |  |  |  |
| Minimum Firing Rate (Hz) | Delayed | 5.3±1.6 | 7.6±2.2*** | 7.3±1.2** | 1.67 | 1.1, 0.3 |
| | Immediate | 5.4±1.6 | 5.7±2.1 | 6.7±2.1 | 0.52 | 10.7, 2.0e-4 |
|  |  | 0.9 |  |  |  | 2.6, 0.09 |
| Maximum Firing Rate (Hz) | Delayed | 46.7±12.4 | 45.4±5.7 | 40.0±5.1 | -0.16 | 1.1, 0.3 |
| | Immediate | 51.2±16.6 | 47.3±14.4 | 49.7±18.9 | -0.52 | 2.2, 0.1 |
|  |  | 0.5 |  |  |  | 1.9, 0.2 |
| Primary Range Gain (Hz/pA) | Delayed | 0.029±0.014 | 0.03±0.19 | 0.023±0.01 | 0.1 | 10.0, 5.1e-3 |
| | Immediate | 0.07±0.04 | 0.07±0.04 | 0.07±0.05 | -0.1 | 0.8, 0.4 |
|  |  | 2.9e-3 |  |  |  | 0.4, 0.7 |
| Sub-primary Range Gain (Hz/pA) | Delayed | 0.07±0.05 | 0.1±0.07 | 0.05±0.02 | 0.66 | 5.8, 0.02 |
|  | Immediate |  |  |  |  |  |

**Table 3:**
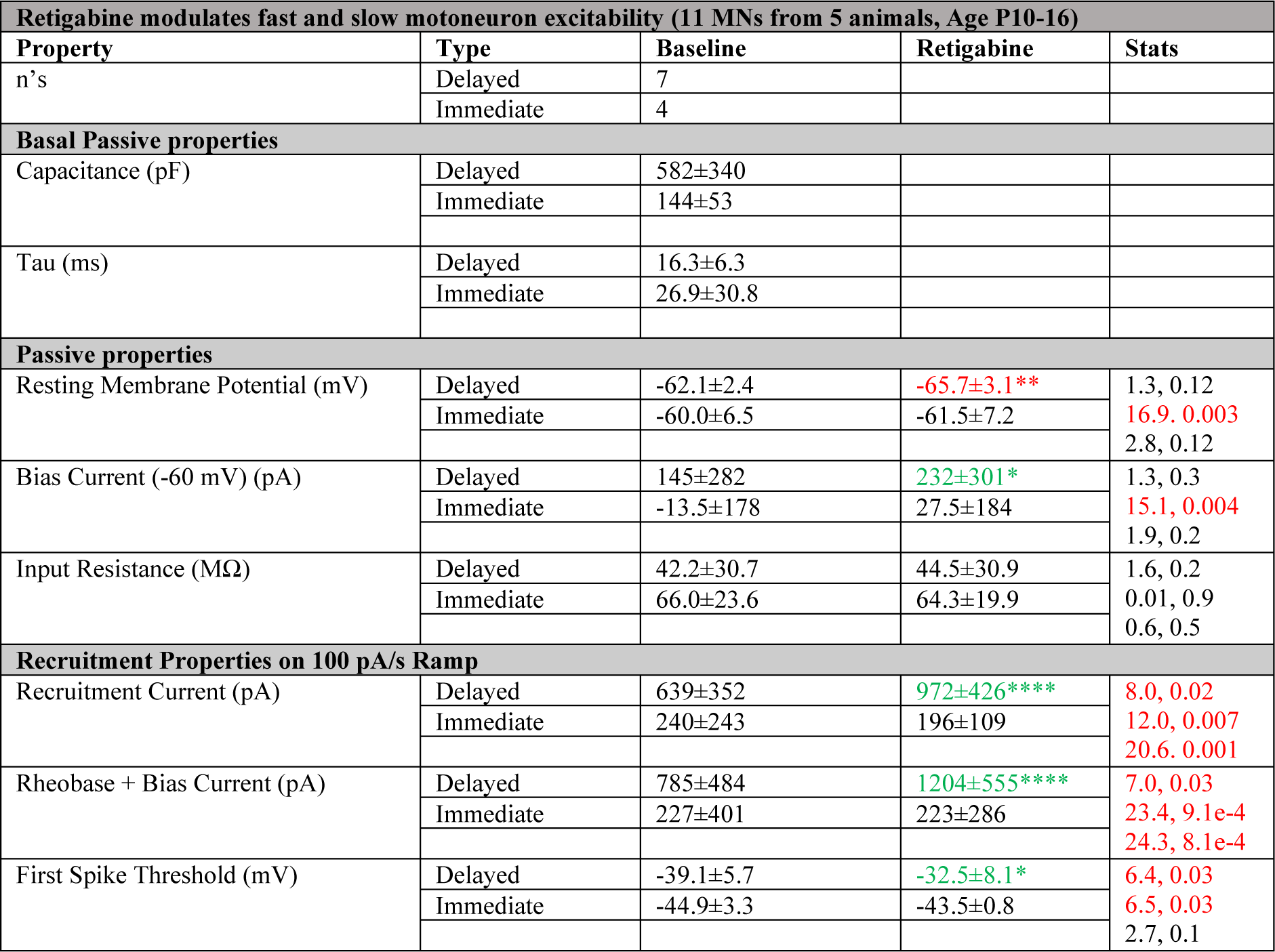
Effects of retigabine on intrinsic properties of motoneuron subtypes. Data are presented as mean ± SD. Asterisks denote significant difference from baseline. F and p values from 2-way Repeated Measures ANOVA for Main effect of MN type, and Main effect of drug, interaction (MN type x Drug). Effect size of agonist: Small: 0.2 – 0.5; medium: 0. 5- 0.8; Large: > 0.8. Asterisks represent differences from baseline p*p<0.05, **p<0.01, ***p<0.001, ****<0.0001.

### M-type potassium currents exert limited control over motoneuron firing gain

Motoneurons produce a non-linear frequency-current (f-I) relationship in response to slow depolarizing current ramps, which can be characterized by a second order polynomial. In addition, we also observed an initial high gain linear region, with variable inter-spike intervals, known as the sub-primary range in delayed firing motoneurons (Figure 4A), which we did not observe in immediate firing motoneurons (Figure 4B). Consistent with the changes in firing threshold reported above, XE991 produced a robust left shift in the f-I relationship of delayed firing motoneurons (left: 11/16; right: 3/16; no change: 2/16; Figure 4A) with few shifts observed in immediate firing motoneurons (left: 2/11; right: 1/11; no change: 8/11; Figure 4B). To examine whether blockade of M-currents influenced firing rate, independent of recruitment, we compared maximum firing rate and slope of the primary and sub-primary ranges before and after administration of XE991. Interestingly, we did not find any changes in maximal firing rate (Figure 4C)**^t^** or f-I gain within the primary range (Figure 4D)**^u^**of either subtype, nor did we find any changes in the gain within the sub-primary range of delayed firing motoneurons (Figure 4E)**^v^**.

As would be expected, activation of KCNQ channels produced a predominant right shift in f-I curves in delayed firing motoneurons (left: 1/11; right: 10/11; Figure 4F) with no consistent shift observed in immediate firing motoneurons (left: 3/10; right: 3/10; no change: 4/10; Figure 4G). However, we found no changes in the maximal firing rate (Figure 4H)**^w^** or primary range gain (Figure 4I)**^x^** in either subtype. Interestingly, in the presence of ICA73, we noted a trend toward an increase in subprimary range gain, which was reversed by XE991 (Figure 4J)**^y^**. Together these data indicate that M-currents exert most of their influence on motoneuron recruitment and produce limited control over firing rate.

### M-current activation decreases monosynaptic reflex recruitment gain

In a final set of experiments, we set out to extrapolate from our single cell studies, in which recruitment was assessed by somatic current injection, to determine if activation of KCNQ channels could produce similar effects on motoneuron recruitment when activated synaptically and assessed at a population level. To address this aim, we performed a study on the monosynaptic reflex (MSR) where the motor pool can be activated synaptically via sensory afferents by delivering electrical stimuli to a dorsal root leading to a short latency (2 ms) population response in the ventral root within the same spinal segment (Figure 5A-C). Consistent with our single cell recordings, activating M-type potassium channels with ICA73 (10 µM) produced a rightward shift in the monosynaptic reflex recruitment curve (Figure 5D), reflected by an increase in MSR threshold (Figure 5E)**^z^**, increase in the half activation current (Figure 5F)**^aa^**, and reduction in the maximum amplitude of the MSR (Figure 5G)**^bb^**. This effect was replicated with a second M-current activator retigabine (10 µM), which also produced a rightward shift in the MSR recruitment curve (Figure 5H), increased MSR threshold (Figure 5I)**^cc^**, half activation current (Figure 5J)^dd^, and reduced maximal MSR amplitude (Figure 5K)**^ee^**.

**Figure 5:**
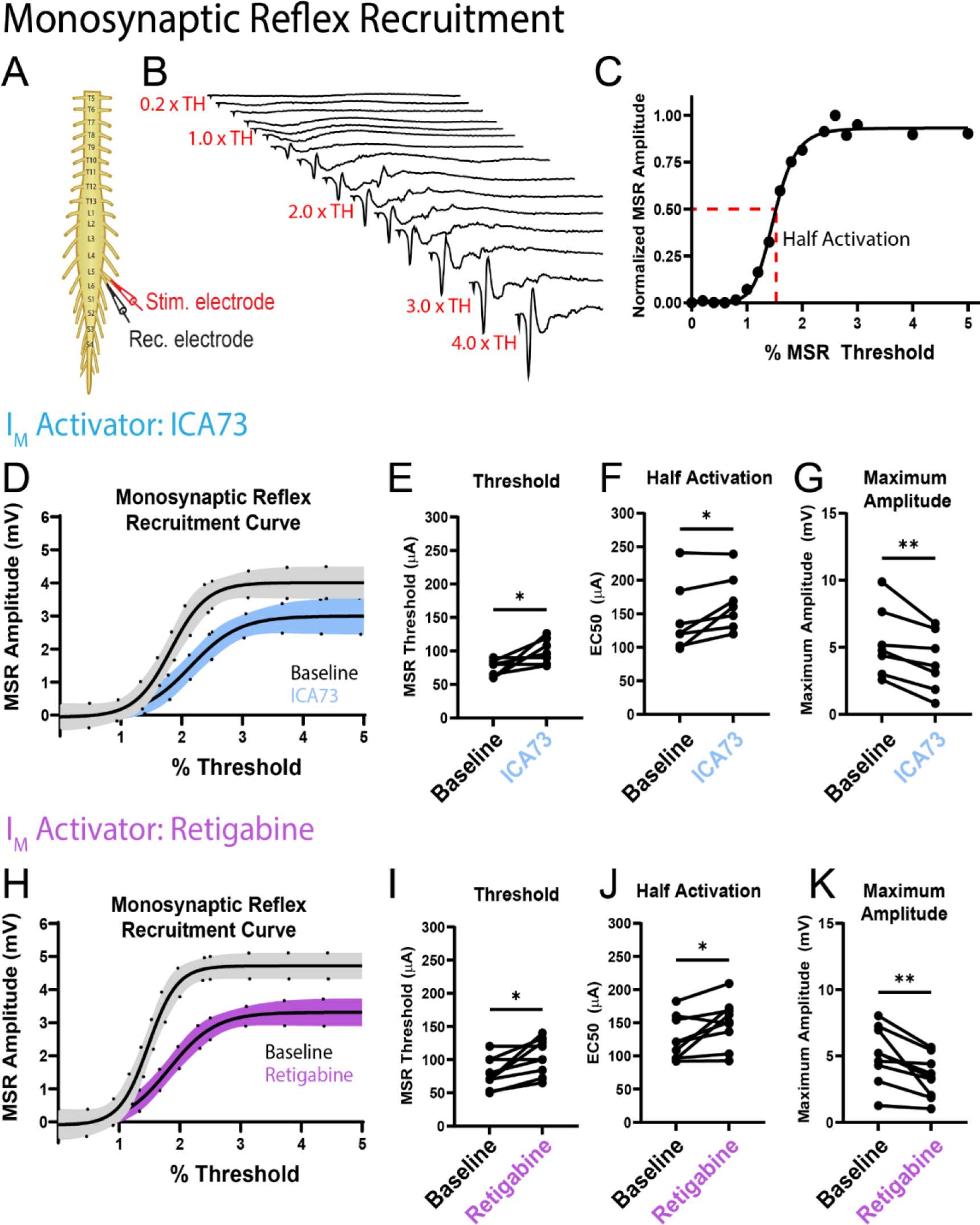
M-currents control motoneuron recruitment on a population level. (A) Isolated spinal cords were obtained from postnatal mice at postnatal days 5-8. Monosynaptic reflexes were recorded in ventral roots in response to brief electrical stimuli applied through a suction electrode to the dorsal root of the fifth lumbar segment (L5). (B &C) MSR recruitment curves were constructed by measuring the peak-to-peak amplitude of the short latency responses that were elicited in response to electrical stimuli that were delivered at 20% increments relative to MSR threshold. MSR recruitment curves were studied before (black) and after application of the KCNQ channel activator ICA73 (blue; n = 7). Activation of KCNQ channels produced a right shift in MSR recruitment curves (D), characterized by an increase in MSR threshold (E), increase in half activation current (F), and decrease in maximal MSR amplitude (G). These effects were recapitulated with a second KCNQ channel activator, retigabine (purple; n = 9), which produced a similar right shift in the MSR recruitment curve (H), characterized by an increase in MSR threshold (I), increase in half activation current (EC50; J), and decrease in maximal MSR amplitude (K). Data are presented as individual data points and analyzed using paired t-tests. MSR recruitment curves are presented as mean MSR amplitude with 95% confidence intervals depicted in grey, purple, or blue, and were analyzed using a 2 factor repeated measures ANOVA. Asterisks denote significance * p <0.05; ** p<0.01.

## Discussion

Recruitment and the subsequent modulation of motor unit firing rates are two key mechanisms for the gradation of muscle force and are important for adjusting movement precision and vigour. Motor unit recruitment was initially suggested to be determined by the passive properties of motoneurons such as their size (Henneman, 1957). However, size is a poor predictor of recruitment order amongst functionally defined motoneurons that innervate different types of muscle fibres (Burke *et al*., 1973; Gustafsson & Pinter, 1984; Cope & Clark, 1991). Our recent work demonstrated a role for active properties, mediated by voltage sensitive ion channels, in adjusting recruitment gain across the motor pool and maintaining orderly recruitment of functionally defined motoneuron subtypes (Sharples & Miles, 2021). In this work, a putative outward current was unmasked in fast and slow motoneurons following blockade of HCN channels. There are several outward currents that are generated by different families of potassium channels that oppose the H-current (Ih) (MacLean *et al*., 2005; Buskila *et al*., 2019). One of these currents is a persistent outward M-type potassium current conducted by KCNQ channels, which not only opposes Ih (MacLean *et al*., 2005; Buskila *et al*., 2019), but also the actions of persistent inward currents (Buskila *et al*., 2019). Here, we demonstrate a role for M-type potassium currents in the regulation of fast but not slow motoneuron recruitment, and a net effect of M-currents on the tuning of recruitment gain across the motor pool. The recruitment gain is defined by the range of input over which a motor pool is recruited. Recruitment of a motor pool over a narrower range of input would be expected to favour the generation of more vigorous movement (Nielsen *et al*., 2019), whilst the recruitment of a motor pool over a broader range of input is likely to favour more precise movements (Figure 6) (Nielsen *et al*., 2019). Our work suggests that M-type potassium currents can adjust the recruitment gain in either direction and may be a target for neuromodulators to readily modify the vigour and precision of movement (Figure 6).

**Figure 6:**
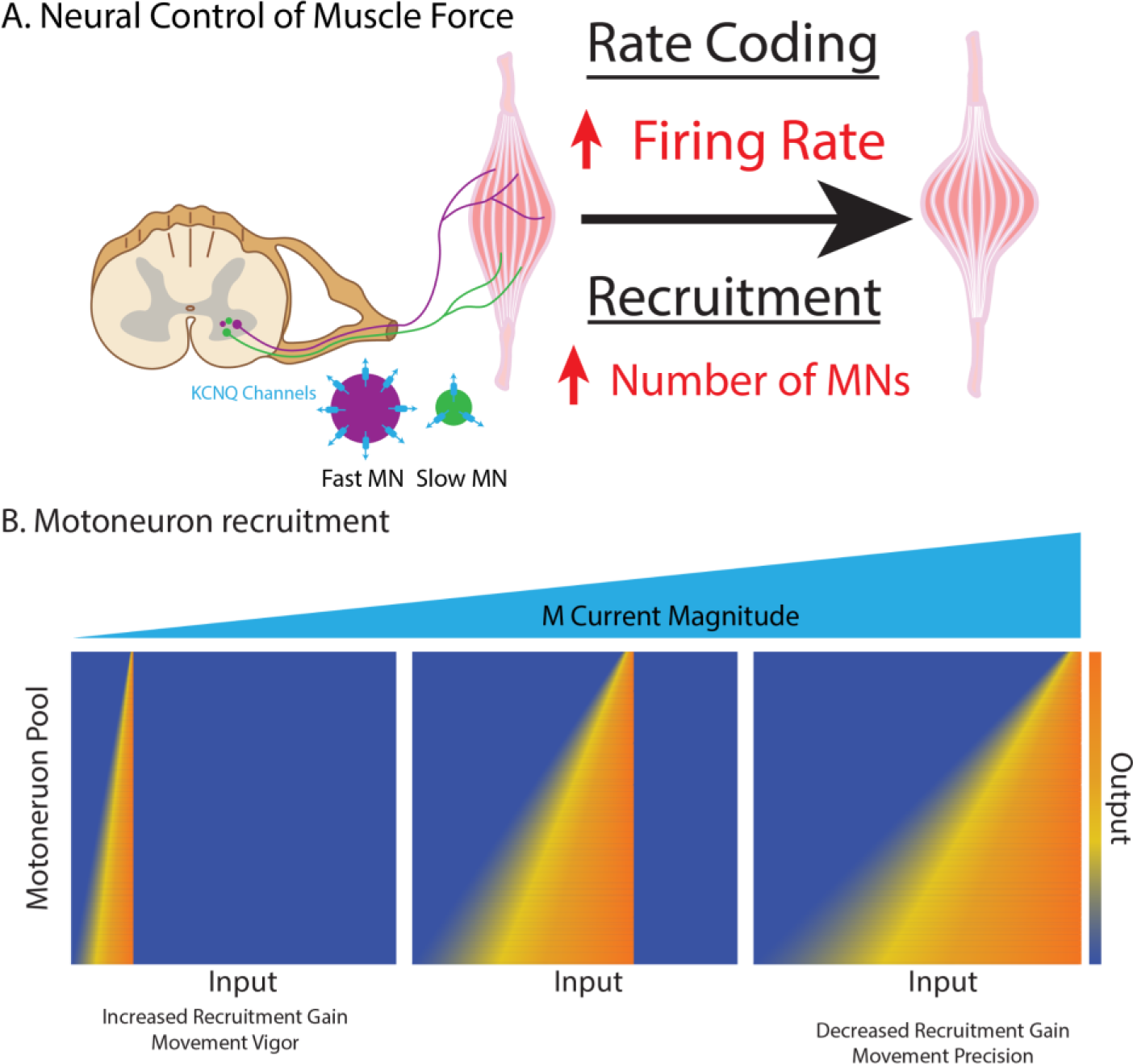
Summary schematic. A.) Recruitment and subsequent modulation of motoneuron firing rates serve as two key mechanisms for the gradation of muscle force. Fast motoneurons (purple) have larger M-currents, conducted by KCNQ channels (blue channels), than slow motoneurons (green). B.) M-Currents (blue triangle) can adjust the range of inputs over which a motor pool is recruited, which may modulate the relative degree of movement precision or vigour.

M-type potassium currents are present in neurons across the central nervous system (Jentsch, 2000) and previous work in motor circuits has described specific roles for M-currents in the control of several intrinsic properties and the firing rates of undefined motoneuron subtypes in the brainstem and lumbar spinal cord (Alaburda *et al*., 2002; Miles *et al*., 2005; Rivera-Arconada & Lopez-Garcia, 2005; Leroy *et al*., 2015; Lombardo & Harrington, 2016; Ghezzi *et al*., 2017, 2018; Lombardo *et al*., 2018). This work has been important in showing that M-currents are present and have a role in motoneuron physiology, however, our work and that of others has shown that recruitment must also be considered along with firing rates, because it’s a key feature that defines motoneuron subtypes and regulates muscle force. This is particularly true given that a large proportion of the M-current is activated within the subthreshold voltage range. In particular, we found that M-currents were larger and had a greater proportion of maximal current activated below spike threshold in fast-like delayed firing motoneurons compared to the slow-like immediate firing motoneurons. Previous work has demonstrated differential expression of KCNQ channel subunits amongst lumbar motoneurons, with all motoneurons expressing Kv7.2 and only half expressing Kv7.3 (Verneuil *et al*., 2020). Given that heteromeric complexes formed between Kv7.2 and 7.3 can produce substantially larger currents than one or the other alone (Wang *et al*., 1998), we hypothesized that differences in current magnitude between subtypes could be accounted for by KCNQ channel subunit expression. However, in contrast to this hypothesis, we found that fast and slow type motoneurons, identified using the molecular markers MMP9 and Errβ, expressed a similarly mixed composition of both Kv7.2 and 7.3 subunits.

Consistent with previous work on undefined motoneuron subtypes in the brainstem and spinal cord (Alaburda *et al*., 2002; Miles *et al*., 2005; Rivera-Arconada & Lopez-Garcia, 2005; Leroy *et al*., 2015; Lombardo & Harrington, 2016; Ghezzi *et al*., 2017), we found that pharmacological manipulation of KCNQ channels produced changes in the intrinsic properties and firing gain; however, these effects were largely restricted to delayed but not immediate firing motoneurons. We reasoned that manipulating M-currents produced selective effects on fast motoneurons because of their larger currents, which are activated mostly below spike threshold. Interestingly, when factoring in changes in firing gain that are due to recruitment, pharmacological manipulation of KCNQ channels produced minimal changes in the firing rate of fast motoneurons. In particular, blocking KCNQ channels with the state-dependent blocker XE991, leads to a robust reduction in the rheobase, which is paralleled by an increase in the input resistance. While XE991 also increased the input resistance of immediate firing motoneurons, this was not sufficient to cause an equivalent change in rheobase. We also found that activation of KCNQ channels with both retigabine and ICA73 increases rheobase, which is paralleled by a depolarization of the spike threshold and a hyperpolarization of the resting potential. Importantly, the effects of ICA73 were reversed by subsequent application of XE991, supporting a role for KCNQ channels in mediating these effects. One possible explanation for the apparent differences is that previous works were conducted on motoneurons at early developmental stages where fast and slow motoneurons are not yet functionally defined (Sharples & Miles, 2021). Alternatively, it is also possible that other studies were biased toward the larger motoneurons, which are more likely to be of the fast subtype.

In our final set of experiments, we set out to extrapolate from our single cell studies, in which recruitment was assessed by somatic current injection, to determine if activation of KCNQ channels could produce similar effects on motoneuron recruitment when activated synaptically and assessed at a population level. In line with our single cell recordings, we found that activation of KCNQ channels with retigabine or ICA73 produced a right shift in monosynaptic recruitment curves, which is consistent with the inhibitory effect that we see on individual motoneurons. In addition to a right shift in the MSR recruitment curve, we also found a robust reduction in the amplitude of the MSR. While MSR amplitude is due to a combination of motoneuron recruitment and firing rate, our single cell data indicates that this inhibitory effect is more heavily weighted by the influence of KCNQ channels on motoneuron recruitment. Given that KCNQ channels are also expressed in DRG neurons and sensory afferents (Devaux *et al*., 2004; Schwarz *et al*., 2006; Wu *et al*., 2017; Zhang *et al*., 2019), it should be noted that changes within sensory pathways could also contribute to the right shift in MSR recruitment curves that we observed.

These results establish roles for voltage gated ion channels in the differential control of motoneuron recruitment, which is important because many of these channels are well known to be targeted by multiple neuromodulatory inputs. Indeed, emerging evidence points toward differential control of motoneuron subtypes, through yet unresolved mechanisms (Bertuzzi & Ampatzis, 2018; Jha & Thirumalai, 2020; Eleftheriadis et al., 2022). M-type potassium currents were initially described based on the inhibitory influence of muscarine on these channels (Brown & Adams, 1980; Wang *et al*., 1998). However, few cholinergic synapses, including C boutons, have been reported along the axon initial segment (AIS) where many of these channels are clustered (Deardorff *et al*., 2021). Although we find KCNQ channel labelling in both fast and slow motoneuron somata, the labelling does not appear to cluster in line with what would be expected at C bouton synapses (Deardorff *et al*., 2013). While cholinergic pathways are likely to modulate M-currents in turtle motoneurons (Alaburda *et al*., 2002), they do not seem to contribute to cholinergic modulation of hypoglossal motoneurons (Ireland *et al*., 2012). Although not tested directly in mouse lumbar motoneurons, pharmacological activation of muscarinic receptors in fast motoneurons produces a very different complement of changes on intrinsic properties compared to manipulating KCNQ channels (Miles *et al*., 2007; Nascimento *et al*., 2019; Eleftheriadis et al., 2022). On the other hand, 5HT fibres have been reported at the AIS (Deardorff *et al*., 2021) where KCNQ channels are clustered (Verneuil *et al*., 2020). In line with these findings, M-currents are reduced by 5HT in neocortical pyramidal (Stephens *et al*., 2018) and hypothalamic neurons (Roepke *et al*., 2012). However, the excitatory actions of 5HT on motoneuron excitability are believed to be largely mediated by the enhancement of persistent inward currents at dendritic synapses (Heckman *et al*., 2003; Perrier *et al*., 2013), with spillover of 5-HT, due to prolonged activation, conversely leading to inhibition of motoneuron output through the activation of extra synaptic receptors on the AIS (Cotel *et al*., 2013). Although structural differences in the AIS have been reported in functionally defined motoneuron subtypes (Rotterman *et al*., 2021), we currently have a limited understanding of whether motoneuron subtypes have different sets of ion channels or neuromodulatory inputs localized to the AIS. The differential neuromodulatory control of motoneuron subtypes therefore serves an exciting direction for future study.

Finally, KCNQ channels have become robust therapeutic targets to combat dysfunction in a range of disorders that are characterized by aberrant neuronal excitability including epilepsy (Gunthorpe *et al*., 2012), chronic pain (Blackburn-Munro & Jensen, 2003), and Amyotrophic Lateral Sclerosis (ALS) (Wainger *et al*., 2014, 2021). While other potassium channels have also been targeted to attenuate neuronal dysfunction in disease (Simon *et al*., 2021), KCNQ channels represent a particularly tractable target given the availability of a growing number of drugs, including those used in clinical trials, that selectively modulate KCNQ channel activity (Liu *et al*., 2021). For example, activation of KCNQ channels can attenuate hallmarks of ALS in both human stem cell-derived motoneurons (Wainger *et al*., 2014) and patients (Wainger *et al*., 2021). Given that fast motoneurons are particularly susceptible to degeneration in ALS (Shaw & Eggett, 2000; Kaplan *et al*., 2014; Martínez-Silva *et al*., 2018), it is particularly important to generate a better understanding of properties that distinguish motoneuron subtypes in order to refine therapeutic approaches. Importantly, our work here suggests that the effects of KCNQ-targeting therapies are likely to be restricted to the susceptible subsets of motoneurons in ALS. In addition, hyperexcitability leading to spasticity is a hallmark of other neurological disorders including but not limited to spinal cord injury (SCI) (Gorassini *et al*., 2004; Elbasiouny *et al*., 2010), cerebral palsy (Bar-On *et al*., 2015; Condliffe *et al*., 2016; Steele *et al*., 2020; Reedich *et al*., 2023), multiple sclerosis (Beard *et al*., 2003), stroke (Udby Blicher & Nielsen, 2009; Burke *et al*., 2013; Mottram *et al*., 2014; Nielsen *et al*., 2019), and dystonia (Pocratsky *et al*., 2023). Although KCNQ channels have not yet been investigated as therapeutic targets in these conditions, they represent promising avenue for future study.

In summary, we demonstrate a role for M-type potassium currents in the differential control of motoneuron subtype recruitment. These data support the notion that motoneuron recruitment is multifaceted and mediated by both active and passive properties. These differences in active properties may therefore provide the capacity to selectively modulate motoneurons in a subtype specific manner or contribute to the differential susceptibility of motoneuron subtypes to degeneration in disease and injury.

## Acknowledgements

The authors reserve the right to apply a Creative Commons Attribution (CC BY) license to any Author Accepted manuscript version arising from this submission. The research data supporting this publication will be made available in an open access data repository upon acceptance to a peer-reviewed journal.

## Funding

This work was supported by fellowships to SAS from the Royal Society (Newton International Fellowship: NIF/R1/180091), Wellcome Trust (ISSF3: 204821/Z/16/Z), and Canadian Institute for Health Research (CIHR-PDF: 202012MFE - 459188 - 297534).

## Contributions

AS, MJB, and JG performed experiments, SAS and MJB analyzed the data, SAS prepared figures, and wrote the manuscript. SAS and GBM conceived and designed the research, interpreted results, and revised the manuscript. All Authors approved the final version of the manuscript.

## Competing Interests

The authors have no conflicting interests to declare.

## Methods

### Animals

Whole-cell patch clamp experiments were performed on tissue obtained from 36 wild type C57 Black J/6 mice of both sexes at postnatal days (P) 10-18. Monosynaptic reflex experiments were performed on 18 animals of both sexes at age P5-8. Immunohistochemistry experiments were performed on spinal cords obtained from animals of both sexes at P12 (n = 4). All procedures were conducted in accordance with the UK Animals (Scientific Procedures) Act 1986, were approved by the University of St Andrews Animal Welfare Ethics Committee, and complies with the animal ethics criteria outlined by the Journal.

### Tissue preparation

All animals were sacrificed by performing a cervical dislocation followed by rapid decapitation. Animals were eviscerated and pinned ventral side up in a dissecting chamber lined with silicone elastomer (Sylguard) filled with carbogen-bubbled (95% oxygen, 5% carbon dioxide) artificial cerebrospinal fluid (aCAF). For monosynaptic reflex experiments, spinal cords were dissected in recording aCSF (containing in mM: 127 NaCl, 3 KCl, 1.25 NaH2PO4, 1 MgCl, 2 CaCl2, 26 NaHCO3, 10 glucose) at room temperature (21 - 23 degrees Celsius). Spinal cords were exposed by performing a ventral vertebrectomy, cutting the ventral roots and gently lifting the spinal cord from the spinal column. Preparations were then transferred to a recording chamber perfused with recirculating, recording aCSF warmed to 25-27 degrees Celsius and given 1 hour recovery time prior to the initiation of baseline measurements.

Preparations used for whole-cell patch clamp electrophysiology were dissected in ice-cold (1-2 degrees Celsius) potassium gluconate-based dissecting/slicing aCSF (containing in mM: 130 K-gluconate, 15 KCl, 0.05 EGTA, 20 HEPES, 25 D-glucose, 3 kynurenic acid, 2 Na-pyruvate, 3 myo-inositol, 1 Na-L-ascorbate; pH 7.4, adjusted with NaOH; osmolarity approximately 345 mOsm). Spinal cords were removed within 3 minutes following cervical dislocation. Spinal cords were secured directly to an agar block (3 % agar) with VetBond surgical glue (3M) and the block glued to the base of the slicing chamber with cyanoacrylate adhesive. The tissue was immersed in ice-cold dissecting/slicing aCSF bubbled with carbogen. Blocks of frozen slicing solution were also placed in the slicing chamber to keep the solution around 1-2 degrees Celsius. On average, the first slice was obtained within 10 minutes of decapitation, which increased the likelihood of obtaining viable motoneurons in slices. 300 µm transverse slices were cut at a speed of 10 um/s on the vibratome (Leica VT1200) to minimize tissue compression during slicing. 3-4 slices were obtained from lumbar segments 4-6 of each animal. Slices were transferred to a recovery chamber filled with pre-warmed (35 degrees Celsius) recovery aCSF (containing in mM: 119 NaCl, 1.9 KCl, 1.2 NaH2PO4, 10 MgSO4, 1 CaCl, 26 NaHCO3, 20 glucose, 1.5 kynurenic acid, 3% dextran), bubbled with carbogen, for thirty minutes after completion of the last slice, which took 10 - 15 minutes on average. Following recovery, slices were transferred to a chamber filled with warm (35 degrees Celsius) recording aCSF and allowed to equilibrate at room temperature (maintained at 23-25 degrees Celsius) for at least one hour before experiments were initiated.

### Whole-cell patch clamp electrophysiology

Whole-cell patch clamp recordings were obtained from 70 lumbar motoneurons. We typically recorded from 2 cells (mode), with a range of 1-4 cells per animal. Slices were stabilized in a recording chamber with fine fibres secured to a platinum harp and visualized with a 40x objective using infrared illumination and differential interference contrast (DIC) microscopy. A large proportion of the motoneurons studied were identified based on location in the ventrolateral region with somata greater than 20 µm. Recordings were also obtained from a subset of motoneurons that had been retrogradely labelled with Fluorogold (Fluorochrome, Denver, CO). Fluorogold was dissolved in sterile saline solution and 0.04 mg/g injected intraperitoneally 24-48 hours prior to experiments (Miles *et al*., 2005). In addition to recording from larger FG-positive cells, this approach allowed us to more confidently target smaller motoneurons. Motoneurons were visualized and whole-cell recordings obtained under DIC using pipettes (L: 100 mm, OD: 1.5 mm, ID: 0.84 mm; World Precision Instruments) pulled on a Flaming Brown micropipette puller (Sutter instruments P97) to a resistance of 2.5-3.5 MΩ. Pipettes were back-filled with intracellular solution (containing in mM: 140 KMeSO4, 10 NaCl, 1 CaCl2, 10 HEPES, 1 EGTA, 3 Mg-ATP and 0.4 GTP-Na2; pH 7.2-7.3, adjusted with KOH).

Signals were amplified and filtered (6 kHz low pass Bessel filter) with a Multiclamp 700B amplifier, acquired at 20 kHz using a Digidata 1440A digitizer with pClamp Version 10.7 software (Molecular Devices) and stored on a computer for offline analysis.

### Identification of fast and slow motoneuron types

Motoneuron subtypes were identified using a protocol established by Leroy et al., (2014, 2015), which differentiates motoneuron type based on the latency to the first spike when injecting a 5 second square depolarizing current. Using this approach, we were able to identify 2 main firing profiles - a delayed repetitive firing profile with accelerating spike frequency, characteristic of fast-type motoneurons, and an immediate firing profile with little change in spike frequency, characteristic of slow-type motoneurons. All motoneuron intrinsic properties were studied by applying a bias current to maintain the membrane potential at -60 mV. Values reported are not liquid junction potential corrected to facilitate comparisons with previously published data (Miles *et al*., 2007; Quinlan *et al*., 2011; Durand *et al*., 2015; Nascimento *et al*., 2019, 2020; Smith & Brownstone, 2020). Cells were excluded from analysis if access resistance was greater than 20 MΩ, changed by more than 5 MΩ over the duration of the recording, or if spike amplitude was less than 60 mV when measured from threshold (described below).

### Data acquisition and analysis

M-currents were measured in voltage clamp mode using an established deactivation protocol (Vernueil et al., 2020). Motoneurons were clamped at a holding potential of -10 mV and subjected to a series of 1 second hyperpolarizing voltage steps (5 mV increments) with a negative deflection in current recorded due to deactivation of KCNQ channels. M-current amplitude was measured as the difference from the initial peak current to steady state current and plotted as a function of voltage.

Basal passive properties including capacitance, membrane time constant (tau), and input resistance (Ri) were measured during a hyperpolarizing current pulse that brought the membrane potential from -60 to -70mV. Input resistance was measured from the initial voltage trough to minimize the impact of active conductances with slow kinetics (eg. Ih, sag). The time constant was measured as the time it took to reach 2/3 of the peak voltage change. Capacitance was calculated by dividing the time constant by the input resistance (C = T/R). Resting membrane potential was measured, from the MultiClamp Commander, 10 minutes after obtaining a whole-cell configuration.

Rheobase and repetitive firing were assessed during slow (100 pA/s) depolarizing current ramps, which allowed us to measure the recruitment current and voltage threshold of the first action potential. Recruitment gain was qualitatively studied by generating cumulative proportion histograms of recruitment current values across all motoneurons samples before and after drug application. The voltage threshold of the first action potential was defined as the voltage at which the change in voltage reached 10 mV/ms. Input resistance was also extracted during the initial linear phase of the subthreshold voltage trajectory during current ramps as in Sharples and Miles (2021). The frequency-current (f-I) relationship during depolarizing current ramps is non-linear. Three main features were extracted from the frequency-current relationship to study firing rate gain - an initial high gain region, with variable interspike interval, known as the sub-primary range, followed by a lower gain, less variable primary firing range, and finally a plateau region where the firing rate reaches a maximum. The slope of the sub-primary range was measured by fitting a line to the initial rise of the f-I relationship. The primary range was captured by extracting the first-rate constant from a second order polynomial function fitted to the f-I relationship (R^2^ = 0.95). Maximal firing rate was defined as the peak frequency reached, which defined the plateau region of the f-I relationship.

### Monosynaptic Reflex

The monosynaptic reflex (MSR) was elicited and recorded with tight fitting suction electrodes attached to the dorsal and ventral roots of the fifth lumbar (L5) segment. Stimuli with a pulse width of 20 us were delivered to the dorsal root at an interval of 60 seconds using a Digitimer DS3 constant current stimulator. In a subset of experiments, we determined that the MSR remained stable over a period of 2 hours and could be reversibly blocked by 2 mM kynurenic acid, which blocks fast glutamatergic synaptic transmission. MSR recruitment curves were constructed by delivering electrical stimuli from 0-5 x MSR threshold intensity (Mean MSR TH: 85± 12 uA) in 20% increments. Recruitment curves were measured before and after a 30-minute drug wash in period. A subset of preparations were then exposed to a wash out with 800 mL of recording aCSF over a period of 60 minutes; however drug effects were not reversed in these preparations. MSR threshold, half max activation intensity (EC50), and maximal amplitude were extracted from recruitment curves for statistical analysis. The EC50 was calculated in Graph Pad using a sigmoidal function fitted to data normalized to maximal MSR amplitude.

### Pharmacology

We deployed pharmacological tools to probe the contribution of ion channels that produce the M-type potassium current. These included the KCNQ channel blocker XE991 (10 uM; Tocris), and two KCNQ channel activators ICA069673 (ICA73; 10 uM; Tocris) and Retigabine (10 uM; Tocris). All drug stocks were prepared in vehicles and concentrations based on vendor recommendations.

### Immunohistochemistry

Neonatal mice were anaesthetised with Pentobarbital then underwent transcardial perfusion with ice-cold phosphate buffered saline (PBS; pH 7.4 without Ca2+ or Mg2+) to remove the blood, followed by ice-cold 4% paraformaldehyde (PFA). The spinal cord was dissected and then incubated in PFA for a further 2-3 hours at 4°C. Spinal cords were then washed in PBS and incubated in 30% sucrose for up to 48 hours. When the spinal cords had sunk in the sucrose solution, they were cryoembedded in Optimal Cutting Temperature compound and frozen at -80°C. Cryosections of 20 µm thickness were obtained and adhered to Superfrost Plus slides (VWR).

Tissue was stored on slides at -80°C. On the day of immunolabelling, slides were thawed and warmed to 37°C in a benchtop oven for 30 mins to aid adherence of the tissue to the slide. Tissue was washed in PBS and then incubated in PBS containing 3% bovine serum albumin (BSA) and 0.2% Triton X100 for 2 hours at room temperature to block non-specific binding and permeabilise the tissue respectively. After blocking/permeabilization, tissue was incubated for 48 hours in PBS containing 1.5% BSA, 0.1% triton X100 and primary antibodies at a dilution of 1 in 500. Primary antibodies are detailed in Table 4. After primary incubation, slides were washed 5 times with PBS over the course of 1 hour. Secondary incubation was performed for 1.5-2 hours in 0.1% Triton X100 with secondary antibodies at final dilutions of 1 in 500. Finally, slides were washed a further 5 times with PBS. Where used, DAPI was diluted to 1 in 2000 in deionised water and samples were incubated for 10 mins at room temperature before being washed in deionised water. Slides were then dried, and coverslips were adhered using Prolong Glass hard-set mountant.

**Table 4:** Antibodies for immunohistochemical analysis of KCNQ channels in motoneuron subtypes.

| Antibody Target | Company | Code | Species | Secondary |
| --- | --- | --- | --- | --- |
| MMP9 | Sigma | M9570 | Goat | Donkey Anti-Goat Alexa Fluor 555 (Abcam, ab150134)<br>Donkey Anti-Goat Alexa Fluor 488 (Invitrogen, A11055) |
| ERR-Beta | Persius Proteomics | PP-H6705-00 | Mouse | Donkey Anti-Mouse Alexa Fluor 405 (Abcam, ab175651),<br>Donkey Anti-Mouse Alexa Fluor 555 (Invitrogen, A31570) |
| CHAT | Milipore | AB144P | Goat | Donkey Anti-Goat Alexa Fluor 555 (Abcam, ab150134) |
| CHAT | Jessell Lab | Lab-Made | Rabbit | Donkey Anti Rabbit Alexa Fluor 647 (Invitrogen, A31573) |
| SMI-32 | Biolegend | 801701 | Mouse | Donkey Anti-Mouse Alexa Fluor 405 (Abcam, ab175651) |
| Kv7.2 | Sigma | Sigma | Rabbit | Donkey Anti Rabbit Alexa Fluor 647 (Abcam, ab150063) |
| Kv7.3 | Alomone | Alomone | Rabbit | Donkey Anti Rabbit Alexa Fluor 647 (Abcam, ab150063) |

### Confocal Microscopy and Image Processing

Confocal microscopy images were captured using a Zeiss LSM800 laser scanning confocal microscope, based on an Axio ‘Observer 7’ microscope, equipped with a 20x 0.8 NA apochromatic objective lens. Illumination was provided by 405, 488, 561 and 640 nm laser lines. Bidirectional laser scanning was performed with 2x line averaging on each channel and images were captured with 8-bit resolution with GaAsP PMT detectors. Images of MMP9 and ERRβ – expressing motoneurons identified using ChAT or SMI32 were captured from 12 serial Z-stacks with XYZ voxel dimensions of 312 x 312 x 600 nm, from which resultant 2D images were produced by summating the intensity across the full z-range. Images of Kv 7.2 and Kv 7.3 expression in either MMP9- or ERRβ-expressing motoneurons were captured using a single-z-plane with resultant XY pixel dimensions of 125 x 125 nm.

Images were visualised and processed using FIJI (Schindelin *et al*., 2012). Analysis of motoneuron subtypes was performed via a semi-automated approach using a custom-designed macro. Briefly, ChAT-or SMI32-positive motoneurons were manually delineated from a smoothed image. MMP9-positive motoneurons were determined based on their intensity of MMP9 expression, whilst ERRβ-positive motoneurons were determined based on the presence of a large binarized nuclear ERRβ structure within the confines of the delineated motoneuron.

### Research design and statistical analysis

Two factor repeated measures ANOVA were conducted to test the effect of pharmacological agents on intrinsic properties and currents, with motoneuron subtype and drug as factors. Appropriate and equivalent nonparametric tests (Mann-Whitney or Kruskal-Wallis) were conducted when data failed tests of normality or equal variance with Shapiro Wilk and Brown-Forsythe tests, respectively. Paired or unpaired t-tests were performed on data with two variables. Individual data points for all cells are presented in figures along with mean ± SD. Statistical analyses were performed using Graph Pad Version 9.0 (Prism, San Diego, CA, USA). Results from all statistical tests performed are reported in Table 5.

**Table 5:** Statistical analyses performed.

|  | Source Data | Condition | Property | N animals | N MNs<br>(Del, Imm) | test | DF | Test<br>statistic | P Value |
| --- | --- | --- | --- | --- | --- | --- | --- | --- | --- |
| **a** | Figure 1, G, J | XE991 | IM Max | 19 | 9,12 | 2WANOVA | 1,19 | 50.2 | 1.0e-6 |
| **b** | Figure 2 D | MN type | IM Max | 25 | 18,15 | T test | 31 | 5.1 | 2.0e-5 |
| **c** | Figure 2 F | MN type | IM ½<br>Activation | 25 | 18,15 | T test | 31 | 0.4 | 0.7 |
| **d** | text | MN type | IM Tau | 25 | 18,15 | T test | 31 | 0.2 | 0.8 |
| **e** | text | Development | IM Amplitude |  | 11,8/ 9,7 | 2WANOVA | 1,31 | 0.04 | 0.8 |
| **f** | Figure 3B | MN type | Spike TH | 25 | 18,15 | T test | 26 | 2.5 | 0.02 |
| **g** | Figure 3C | MN type | IM at Spike<br>TH | 25 | 18,15 | T test | 32 | 4.7 | 4.5e-5 |
| **h** | Figure 3F | XE991 | Rheobase | 17 | 16,13 | 2WANOVA | 1,27 | 9.0 | 0.006 |
| **i** | Figure 3 H | XE991 | Input Res. | 17 | 16,13 | 2WANOVA | 1,27 | 11.6 | 0.002 |
| **j** | Figure 3I | XE991 | Spike TH | 17 | 16,13 | 2WANOVA | 1,27 | 0.8 | 0.4 |
| **k** | Text | XE991 | RMP | 17 | 16,13 | 2WANOVA | 1,27 | 0.1 | 0.7 |
| **l** | Text | XE991 | Ibias | 17 | 16,13 | 2WANOVA | 1,27 | 1.2 | 0.3 |
| **m** | Figure 3L | ICA73 | Rheobase | 14 | 11,11 | 2WANOVA | 2,40 | 7.7 | 0.001 |
| **n** | Figure 3N | ICA73 | Input Res. | 14 | 11,11 | 2WANOVA | 2,40 | 0.4 | 0.6 |
| **o** | Figure 3O | ICA73 | Spike TH | 14 | 11,11 | 2WANOVA | 2,40 | 6.1 | 0.005 |
| **p** | Text | ICA73 | RMP | 14 | 11,11 | 2WANOVA | 2,40 | 4.4 | 0.02 |
| **q** | Table 3 | Retigabine | Rheobase | 5 | 7,4 | 2WANOVA | 1,9 | 20.6 | 0.001 |
| **r** | Table 3 | Retigabine | Spike TH | 5 | 7,4 | 2WANOVA | 1,9 | 6.5 | 0.03 |
| **s** | Table 3 | Retigabine | RMP | 5 | 7,4 | 2WANOVA | 1,9 | 16.7 | 0.003 |
| **t** | Figure 4C | XE991 | Max FR | 17 | 16,13 | 2WANOVA | 1,26 | 0.08 | 0.8 |
| **u** | Figure 4D | XE991 | PR Gain | 17 | 16,13 | 2WANOVA | 1,26 | 1.1 | 0.3 |
| **v** | Figure 4E | XE991 | SPR Gain | 17 | 16,13 | T test | 15 | 1.2 | 0.3 |
| **w** | Figure 4H | ICA73 | Max FR | 14 | 11,11 | 2WANOVA | 2,40 | 1.9 | 0.2 |
| **x** | Figure 4I | ICA73 | PR Gain | 14 | 11,11 | 2WANOVA | 2,40 | 0.4 | 0.7 |
| **y** | Figure 4J | ICA73 | SPR Gain | 14 | 11,11 | RMANOV<br>A | 2,20 | 5.8 | 0.02 |
| **z** | Figure 5E | ICA73 | MSR TH | 7 | 7 | T test | 6 | 2.7 | 0.04 |
| **aa** | Figure 5F | ICA73 | MSR ½ Act. | 7 | 7 | T test | 6 | 2.7 | 0.04 |
| **bb** | Figure 5G | ICA73 | MSR Max | 7 | 7 | T test | 6 | 4.2 | 0.006 |
| **cc** | Figure 5I | Retigabine | MSR TH | 9 | 9 | T test | 8 | 3.2 | 0.01 |
| **dd** | Figure 5J | Retigabine | MSR ½ Act. | 9 | 9 | T test | 8 | 2.6 | 0.03 |
| **ee** | Figure 5K | Retigabine | MSR Max | 9 | 9 | T test | 8 | 3.6 | 0.007 |

## Notes

### Competing Interest Statement

The authors have declared no competing interest.

